# *Medicago truncatula* copper transporter 1 (MtCOPT1) delivers copper for symbiotic nitrogen fixation

**DOI:** 10.1101/104554

**Authors:** Marta Senovilla, Rosario Castro-Rodríguez, Isidro Abreu, Viviana Escudero, Igor Kryvoruchko, Michael K. Udvardi, Juan Imperial, Manuel González Guerrero

**Author notes:** Present address: Bioengineering Department, Kafkas University, Kars 36100, Turkey. Corresponding author: Manuel González-Guerrero.

## Abstract

• Copper is an essential nutrient for symbiotic nitrogen fixation. This element is delivered by the host plant to the nodule, where membrane copper transporter would introduce it into the cell to synthesize cupro-proteins.

• COPT family members in model legume *Medicago truncatula* were identified and their expression determined. Yeast complementation assays, confocal microscopy, and phenotypical characterization of a *Tnt1* insertional mutant line were carried out in the nodule-specific *M.truncatula* COPT family member.

• *Medicago truncatula* genome encodes eight COPT transporters. *MtCOPT1* (*Medtr4g019870*) is the only nodule-specific *COPT* gene. It is located in the plasma membrane of the differentiation, interzone and early fixation zones. Loss of MtCOPT1 function results in a copper-mitigated reduction of biomass production when the plant obtains its nitrogen exclusively from symbiotic nitrogen fixation. Mutation of *MtCOPT1* results in diminished nitrogenase activity in nodules, likely an indirect effect from the loss of a copper-dependent function, such as cytochrome oxidase activity in *copt1-1* bacteroids.

• These data are consistent with a model in which MtCOPT1 transports copper from the apoplast into nodule cells to provide copper for essential metabolic processes associated with symbiotic nitrogen fixation.

## Introduction

Symbiotic nitrogen fixation is the conversion of N_2_ into NH_4_^+^ carried out by bacteria associated with host organisms, such as rhizobia inside legume root nodules (van Rhijn & Vanderleyden, 1995; Oldroyd, 2013; Downie, 2014). Legume root nodules result from a complex developmental program initiated by the exchange of chemical signals between the symbionts (Oldroyd, 2013; Antolín-Llovera et al., 2014). During this process, cells from the root cortex and from the endodermis and pericycle proliferate to form the nodule primordium (Xiao et al., 2014). In parallel, rhizobia are guided by infection threads from the root hairs to the inner cell layers of the nodule. There, in an endocytic-like process, they are released into the host cell cytosol, surrounded by a plant membrane called the symbiosome membrane (SM) (Limpens et al., 2009). The SM and enclosed rhizobia constitute a specialized, albeit transient, organelle called the symbiosome. Rhizobia within symbiosomes divide and eventually differentiate into nitrogen-fixing bacteroids as a microaerobic environment is established in the developing nodule (Vasse et al., 1990; Miller et al., 1993; Bobik et al., 2006). Two different nodule developmental programs are known: indeterminate, as in *Medicago* and *Pisum*; and determinate, as in *Lotus* and *Glycine*. The main difference is the persistence of a nodule meristem(s) in the indeterminate type, which gives rise to four contemporaneous developmental zones: the meristem (zone I); the infection/differentiation region where rhizobia are released from infection threads and differentiate into bacteroids (zone II) the nitrogen fixation zone (III); and the senescent zone (IV) (Vasse et al., 1990). In addition to this, some authors propose additional regions such as the interzone between Zone II and Zone III (Roux et al., 2014).

Transition metals, such as iron and copper, play an important role in symbiotic nitrogen fixation (Brear et al., 2013; González-Guerrero et al., 2014; González-Guerrero et al., 2016). Reduced levels of these metals in plants, caused by the low bioavailability of these nutrients in some soil types, have a detrimental effect on nitrogen fixation rates (Tang et al., 1992; Ibrikci & Moraghan, 1993; O'Hara, 2001). This results from their role as cofactors of many of the key enzymes involved symbiotic nitrogen fixation. Iron is part of the heme group that allows leghemoglobin to bind, transport, and buffer the nodule free O_2_ at nanomolar levels and, thus, avoid nitrogenase poisoning (Appleby, 1984; Ott et al., 2005). Iron is also at the catalytical core of enzymes involved in free radical control (Hersleth et al., 2006) and in the metallic clusters in nitrogenase (Miller et al., 1993; Rubio & Ludden, 2005). Copper is used in free radical metabolism in the nodule, as part of Cu,Zn superoxide dismutase (Rubio et al., 2004). It is also a cofactor of cytochrome oxidase, the final complex of an electron transport chain that reduces oxygen for energy metabolism. This system is essential for rhizobial survival, given their strictly aerobic metabolism. In fact, to survive in the microaerobic environment in the nodule and to satisfy the high-energy demands of symbiotic nitrogen fixation, a high-affinity copper-containing cytochrome *cbb*_*3*_ oxidase is expressed by the bacteroids (Soupène et al., 1995; Preisig et al., 1996b; Udvardi & Poole, 2013). Loss of the activity of this enzyme results in loss of nitrogen fixation (Preisig et al., 1993), very likely as a consequence of the inability to provide sufficient energy to maintain nitrogenase activity at high enough levels, or of the increased oxygen levels that might result from missing this high-affinity O_2_ reducing system.

Metallic micronutrients in rhizobia have to be provided by the host plant (Johnston et al., 2001). In the case of iron, and likely also for copper, they are carried by the vasculature from the root and released in the apoplast of the infection/differentiation zone in indeterminate type nodules, such as those of *Medicago truncatula* (Rodríguez-Haas et al., 2013). From there, a number of metal transporters introduce them in the cytosol of rhizobia-infected cells. In the case of iron, this is mediated by MtNramp1 (Tejada-Jiménez et al., 2015). In the case of copper, this could be mediated by a YSL (Yellow Stripe-Like) transporter if the substrate is nicotianamine-bound metal (Schaaf et al., 2004; Conte & Walker, 2011; Zheng et al., 2012), or by a COPT (Copper transporter) protein if it is Cu^+^ (Sancenón et al., 2003; Pilon, 2011). Although nicotianamine seems to be important for SNF, as indicated by the loss of fixation capabilities in a mutant affected in the capability to synthesize this chelator (Avenhaus et al., 2016), the fact that iron is incorporated as Fe^2+^ (since it is mediated by a Nramp transporter) (Tejada-Jiménez et al., 2015) suggests that nicotianamine, and hence YSL proteins, is not be directly involved in metal uptake by rhizobia-infected cells.

COPT transporters, also known as Ctr in fungi and animals, are present in all eukaryotes (Lee et al., 2001; Zhou & Thiele, 2001; Sancenón et al., 2003). They are trimeric proteins in which the monomer is a 140-400 amino acid polypeptide with three transmembrane regions (Dumay et al., 2006). This protein has methionine-rich regions frequently present in the N-terminal extracytosolic region and a conserved MXXXM motif in the second transmembrane domain (De Feo et al., 2009). The later motif forms a channel-like structure in the trimer that binds two Cu^+^ ions/trimer in a trigonal planar way (Dumay et al., 2006). In plants, COPT proteins constitute multigenic families (Sancenón et al., 2003; Yuan et al., 2011). Some of their members appear to be responsible for copper uptake from soil (Sancenon et al., 2004), in a manner very similar to that of iron uptake in Strategy I plants, *i.e.* transport of reduced metal produced by a ferroreductase-oxidase (FRO) protein, either FRO4 or FRO5 in *Arabidopsis thaliana* (Bernal et al., 2012). These transporters have also been associated with pollen development (Sancenon et al., 2004), remobilization of stored copper (Garcia-Molina et al., 2011), and copper allocation in aerial tissues (Garcia-Molina et al., 2013). However, in spite of the importance of copper in symbiotic nitrogen fixation (O'Hara, 2001; González-Guerrero et al., 2014; González-Guerrero et al., 2016), and the role of COPT proteins in copper uptake (Puig et al., 2007; Pilon, 2011), very little information is available about the role of these proteins in the nodule.

In this study, we have identified *MtCOPT1* (*Medtr4g019870*) a gene encoding a nodule-specific COPT family member. Consistent with a role in copper uptake from the apoplast, *MtCOPT1* expression is mainly confined to the differentiation/interzone region of the nodule, where its encoded protein is located in the plasma membrane of those cells. Characterization of a *Tnt1* insertional mutant in this gene showed a reduction in plant biomass production during symbiosis, associated with decreases in bacteroid cytochrome oxidase and nitrogenase activities. This was restored by transformation with a wild-type copy of the mutated gene or by increasing copper concentrations in the nutrient solution. This work adds to our understanding of how copper is delivered from the host plant to be used in symbiotic nitrogen fixation.

## Materials and Methods

### Biological materials and growth conditions

*Medicago truncatula* R108 seeds were scarified in concentrated H_2_SO_4_ for 7 min. Then, they were washed with cold water and surface-sterilized with 50% bleach for 90 s and incubated overnight in sterile water for imbibition. After 48 h at 4ºC, seeds were germinated in water-agar plates at 22ºC for 24 h. Then, seedlings were planted in sterile perlite pots and inoculated with *Sinorhizobium meliloti* 2011 or *S. meliloti* 2011 transformed with pHC60 (Cheng & Walker, 1998), as indicated. Plants were cultivated in a greenhouse in 16 h of light and 22ºC conditions, and watered every two days with Jenner’s solution or water, alternatively (Brito et al., 1994). Nodules were collected at 28 dpi. Non-nodulated plants were grown in similar conditions of light and temperature but instead of being inoculated with *S. meliloti,* they were watered every two weeks with solutions supplemented with 2 mM NH_4_NO_3_. For hairy-root transformations, *M.truncatula* seedlings were transformed with *Agrobacterium rhizogenes* ARqua1 carrying the appropriate binary vector as described (Boisson-Dernier et al., 2001). In agroinfiltration experiments, tobacco (*N. benthamiana*) leaves were transformed with the plasmid constructs in *A. tumefaciens* C58C1 (Deblaere et al., 1985). Tobacco plants were grown in a greenhouse under the same conditions as *M. truncatula*.

*Saccharomyces cerevisiae* strain *Δctr1* and its parental strain BY4741 (MATa *his3Δ1 leu2Δ0 met15Δ0 ura3Δ0*) were purchased from the Yeast Knockout Collection (GE Dharmacon) and used for heterologous expression assays. Yeasts were grown in synthetic dextrose (SD), or in yeast peptone dextrose (YPD) media supplemented with 2% glucose (Sherman et al., 1986). Phenotypic characterization was done in yeast peptone ethanol glycerol (YPEG) medium (Li & Kaplan, 2001).

### Quantitative real-time RT-PCR

Transcriptional expression studies were carried out by real-time RT-PCR (StepOne plus, Applied Biosystems) using the Power SyBR Green master mix (Applied Biosystems). Primers used are indicated in Supplemental Table 1. RNA levels were normalized by using the *ubiquitin carboxy-terminal hydrolase* gene as internal standard for *M. truncatula* genes, and *pyruvate dehydrogenase B* for *S. meliloti* transcripts. RNA isolation and cDNA synthesis were carried out as previously described (Tejada-Jiménez et al., 2015).

### Yeast complementation assays

*MtCOPT1* cDNA was cloned between the *Xba*I and *Eco*RI sites of the expression vector pYPGE15. Restriction sites were added to *MtCOPT1* CDS by PCR, using the primers listed (Supporting Information, Table S1). Yeast were transformed using a lithium acetate-based method (Schiestl & Gietz, 1989). Transformants were selected in SD medium by uracil autotrophy. For phenotypic tests, transformants were plated in YPEG (Li & Kaplan, 2001) supplemented or not with 1 mM CuSO_4_.

### GUS staining

Two kb upstream of *MtCOPT1* start codon were amplified using the primers indicated in Supporting Information, Table S1, then cloned in pDONR207 (Invitrogen) and transferred to destination vector pGWB3 (Nakagawa et al., 2007) using Gateway Cloning technology (Invitrogen). An *A. rhizogenes* ARqua1 derived strain transformed with this pGWB3-based vector was used for hairy root transformation of *M. truncatula* plants as indicated (Boisson-Dernier et al., 2001). Transformed plants were transferred to sterilized perlite pots and inoculated with *S. meliloti* 2011. GUS activity was determined in 28 dpi plants as described (Vernoud et al., 1999).

### Immunohistochemistry and confocal microscopy

A DNA fragment integrating the full length *MtCOPT1* genomic region and the two kb upstream of its start codon with three HA epitopes fused in N-terminus of the protein, was cloned in the plasmid pGWB1 (Nakagawa et al., 2007) using Gateway technology (Invitrogen). In frame fusion of the epitopes was done by fusion PCR using the primers indicated in Supporting Information, Table S1. Hairy-root transformation was performed as previously described (Boisson-Dernier et al., 2001). Transformed plants were transferred to sterilized perlite pots and inoculated with *S. meliloti* 2011 containing the pHC60 plasmid that constitutively expresses GFP. Nodules collected from 28-dpi plants were fixed by overnight incubation in 4% paraformaldehyde, 2.5% sucrose in PBS at 4ºC. After washing in PBS, nodules were cut in 100 μm sections with a Vibratome 1000 plus (Vibratome). Sections were dehydrated using methanol series (30, 50, 70, 100% in PBS) for 5 min and then rehydrated. Cell walls were permeabilized with 4% cellulase in PBS for 1 h at room temperature and with 0.1% Tween 20 in PBS for 15 min. Sections were blocked with 5 % bovine serum albumin (BSA) in PBS before their incubation with an anti-HA mouse monoclonal antibody (Sigma) for 2 hours at room temperature. After washing, an Alexa594-conjugated anti-mouse rabbit monoclonal antibody (Sigma) was added to the sections for 1 h at room temperature. DNA was stained with DAPI after washing. Images were acquired with a confocal laser-scanning microscope (Leica SP8) using excitation lights at 488 nm for GFP and at 561 nm for Alexa 594.

### Transient *MtCOPT1* expression in Tobacco leaves

*MtCOPT1* coding sequence was fused to GFP at N-terminus by cloning in pGWB6 (Nakagawa et al., 2007) using Gateway Technology (Invitrogen). These constructs, and the plasma membrane marker pm-CFP pBIN (Nelson et al., 2007) were introduced into *A. tumefaciens* C58C1 (Deblaere et al., 1985). Transformants were grown in liquid medium to late exponential phase. Then, cells were centrifuged and resuspended to an OD_600_ of 1.0 in 10 mM MES pH 5.6, containing 10 mM MgCl_2_ and 150 μM acetosyringone. These cells were mixed with an equal volume of *A. tumefaciens* C58C1 expressing the silencing suppressor p19 of Tomato bushy stunt virus (pCH32 35S:p19) (Voinnet et al., 2003). Bacterial suspensions were incubated for 3 h at room temperature and then injected into young leaves of 4 week-old *Nicotiana benthamiana* plants. Leaves were examined after 3 days by confocal laser-scanning microscopy (Leica SP8) with excitation lights of 405 nm for CFP and 488 nm for GFP.

### Acetylene reduction assay

Nitrogenase activity was measured by the acetylene reduction assay (Hardy et al., 1968). Nitrogen fixation was assayed in mutant and control plants at 28 dpi in 30 ml vials fitted with rubber stoppers. Each vial contained four or five pooled transformed plants. Three ml of air inside of the vial was replaced with 3 ml of acetylene. Tubes were incubated at room temperature for 30 min. Gas samples (0.5 ml) were analyzed in a Shimadzu GC-8A gas chromatograph fitted with a Porapak N column. The amount of ethylene produced was determined by measuring the height of the ethylene peak relative to background. Each point consists of two vials each. After measurements, nodules were recovered from roots to measure their weight.

### Metal content determination

Total reflection X-ray fluorescence (TXRF) analysis was used to determine copper content in three sets of 28 dpi nodules, each set originating from the nodules pooled from five plants. Analyses were carried out at Total Reflection X-Ray Fluorescence laboratory from Interdepartmental Research Service (SIdI), Universidad Autónoma de Madrid (Spain). Inductively coupled plasma mass spectrometry (ICP-MS) was carried out in three sets of 28 dpi roots and shoots, each set originating from the nodules pooled from five plants. ICP-MS was carried out at the Unit of Metal Analysis from the Scientific and Technology Centre, Universidad de Barcelona (Spain).

### Cytochrome oxidase activity

Nodules from 28-dpi plants were excised from the root and used for bacteroid isolation, as described by Brito et al. (1994) with modifications. Final resuspension was performed in 50 mM 4-(2-hydroxyethyl)-1-piperazineethanesulfonic acid (HEPES), 200 mM NaCl pH 7.0. Cytochrome oxidase activity was assessed using N,N,N’,N’-tetramethyl-p-phenylenediamine (TMPD) oxidation assay. The reaction was started by adding TMPD to the bacteroid suspension to a final concentration of 2.7 mM. Each sample was measured at OD_520_ each 10 s for 5 min to determine the reaction kinetics. To measure protein content to calculate specific activity, the bacteroid suspension was lysed in 10 % SDS at 90ºC for 5 min. Protein content was measured with the Pierce™ BCA Protein Assay (Thermo Scientific), incubated for 30 minutes at 37 ºC and the absorbance estimated at OD_562_.

### Bioinformatics

To identify *M. truncatula* COPT family members, BLASTN and BLASTX searches were carried out in the *M. truncatula* Genome Project site (http://www.jcvi.org/medicago/index.php). Sequences from model COPT proteins were obtained from JCVI (http://www.jcvi.org/medicago/index.php), TAIR (https://www.arabidopsis.org/), Rice Genome Annotation Project (http://rice.plantbiology.msu.edu) and Uniprot (http://www.uniprot.org): *M. truncatula* (MtCOPT1 to MtCOPT8: Medtr4g019870, Medtr7g066070, Medtr3g105330, Medtr4g064963, Medtr4g065660, Medtr1g015000, Medtr4g065123, Medtr0027s0220), *Arabidopsis thaliana* (AtCOPT1 to AtCOPT6: At5g59030, At3g46900, At5g59040, At2g37925, At5g20650, At2g26975), *Oryza sativa* (OsCOPT1 to OsCOPT7: Os01g56420, Os01g56430, Os03g25470, Os04g33900, Os05g35050, Os08g35490, Os09g26900), *Brachypodium distachion* (BdCOPT1 to BdCOPT5: Bradi1g24180, Bradi1g24190, Bradi2g51210, Bradi4g31330, Bradi5g09580), *Glycine max* (GmCOPT1 to GmCOPT9: Glyma_11g134700, Glyma_18g191300, Glyma_04g057000, Glyma_06g057400, Glyma_01g106700, Glyma_07g141200, Glyma_07g141600, Glyma_14g107100, Glyma_17g219400, Glyma_18g191900), *Phaseolus vulgaris* (PvCOPT1 to PvCOPT6: Phavu_011g060400g, Phavu_011g060500g, Phavu_008g112800g, Phavu_009g083400g, Phavu_008g113200g, Phavu_009g083400g), *Solanum lycopersicum* (SlCOPT1 to SlCOPT8: Solyc02g082080, Solyc09g014870, Solyc06g005820, Solyc08g006250, Solyc10g084980, Solyc01g107640, Solyc09g011700, Solyc06g005620) and *Populus trichocarpa* (PtCOPT1 to PtCOPT9: Poptr_0009s04370g, Poptr_0001s25290g, Poptr_0009s04360g, Poptr_0006s09440g, Poptr_0006s23580g, Poptr_0006s14310g, Poptr_0006s09430g). Trees were constructed from a ClustalW multiple alignment of the sequences (http://www.ebi.ac.uk/Tools/msa/clustalw2), then analyzed by MEGA7 (Tamura *et al.,* 2013) using a Neighbour-Joining algorithm with bootstrapping (1,000 iterations). Unrooted trees were visualized with FigTree (http://tree.bio.ed.ac.uk/software/figtree).

### Statistical tests

Data were analyzed with Student’s unpaired t test to calculate statistical significance of observed differences. Test results with p-values lower than 0.05 were considered as statistically significant.

## Results

### *MtCOPT1* is specifically expressed in nodules

The M. truncatula genome encodes eight COPT genes (*MtCOPT1, Medtr4g019870; MtCOPT2, Medtr7g066070; MtCOPT3, Medtr3g105330; MtCOPT4, Medtr4g064963; MtCOPT5, Medtr4g065660; MtCOPT6, Medtr1g015000; MtCOPT7, Medtr4g065123; and MtCOPT8, Medtr0027s0220*). Their expression profiles were determined in shoots, roots, and nodules in nodulated and non-nodulated plants. Out of the eight, *MtCOPT1* was the only gene whose transcripts were detected in nodules exclusively (Fig. 1a). *MtCOPT3* was the only other *COPT* gene also expressed in nodules, but it was not specific to this organ and its maximum expression occured in shoots, regardless of the symbiotic status of the plant (Supporting Information, Fig. S1). *MtCOPT4* and *MtCOPT6* transcripts were detected only in shoots of nodulated and non-nodulated plants, while *MtCOPT5* was mostly confined to roots. *MtCOPT8* transcripts were detected at low levels in both shoots and roots. No expression of *MtCOPT2* or *MtCOPT7* was observed in any of the samples assessed (Supporting Information, Fig. S1).

**Figure 1.**
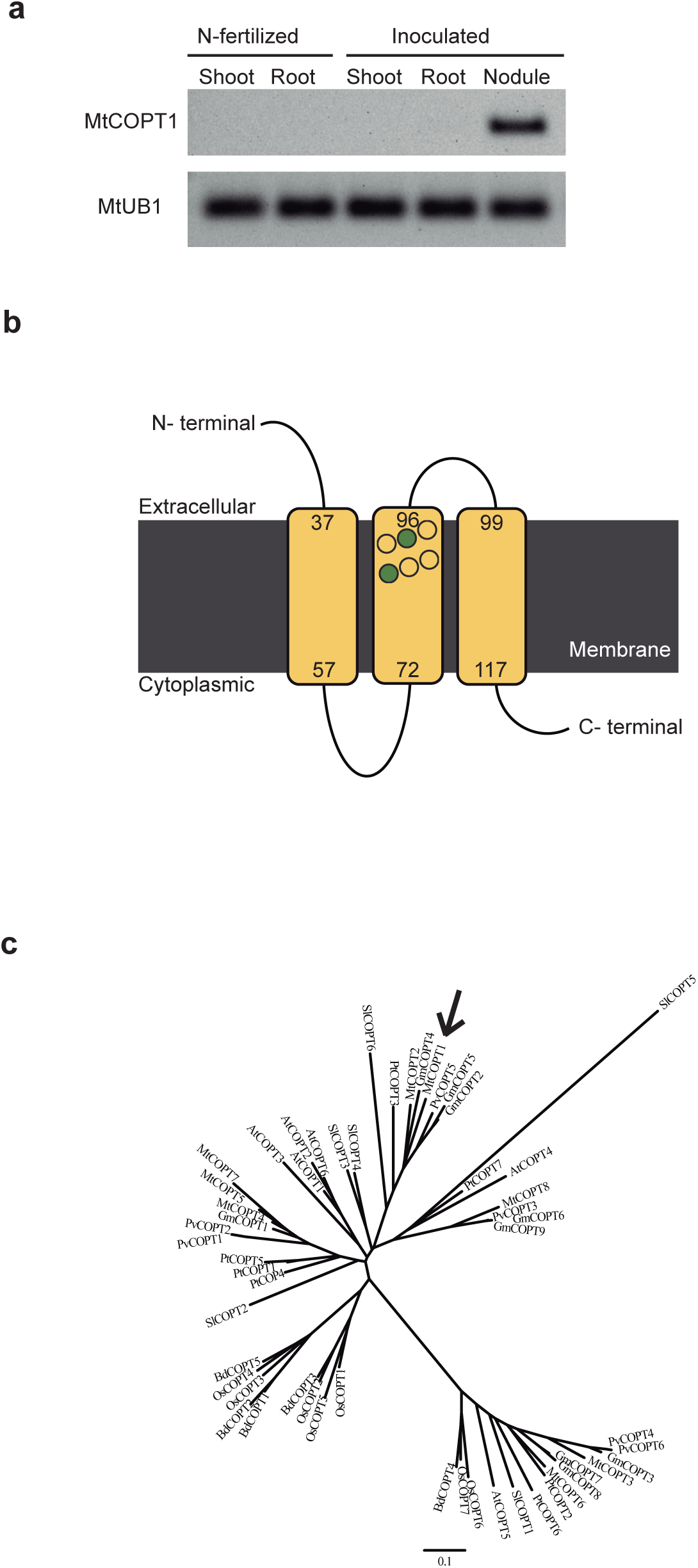
*MtCOPT1* is a member of the COPT gene family and it is specifically expressedin nodules. (a) *MtCOPT1* expression in nodulated and non-nodulated plants. *M.truncatula ubiquitin carboxy-terminal hydrolase1* (MtUB1) expression was used aspositive control for RT-PCR. (b) Proposed topology of MtCOPT1. The conserved MXXXM motif is marked with circles, with methionines indicated in green. (c) Unrooted tree of *M. truncatula* COPT transporters, and representative plant COPT homologues.

Sequence analyses of MtCOPT1 showed the conserved features of COPT1 proteins (Dumay et al., 2006; De Feo et al., 2009), with three predicted transmembrane domains, and a conserved MXXXM domain in the second transmembrane region (Figure 1b). Sequence comparison of *M. truncatula* COPT transporters with those from other sequenced dicots and monocots revealed two major clusters of related sequences (Fig. 1c). MtCOPT1 is located in a branch shared by a subset of legume COPT proteins.

### MtCOPT1 transports copper towards the cytosol

To confirm that MtCOPT1 was able to transport copper, yeast complementation assays were carried out using a *S. cerevisiae* mutant with a deletion in the *ScCTR1* gene. This mutant is affected in Cu^+^ uptake and, consequently, copper dependent metabolic reactions are affected. For instance, this strain is not able to grow on non-fermentable carbon sources since it lacks the copper-dependent cytochrome oxidase activity required. Therefore, this mutant was not able to grow on YPEG medium, that contains ethanol and glycerol as carbon sources (Li & Kaplan, 2001), unless the copper concentration of the medium was increased (Fig. 2). However, when these mutants were transformed with a vector expressing *MtCOPT1,* growth on YPEG under low copper conditions was restored to almost wild-type levels, indicating a role of MtCOPT1 in Cu^+^ uptake.

**Figure 2.**
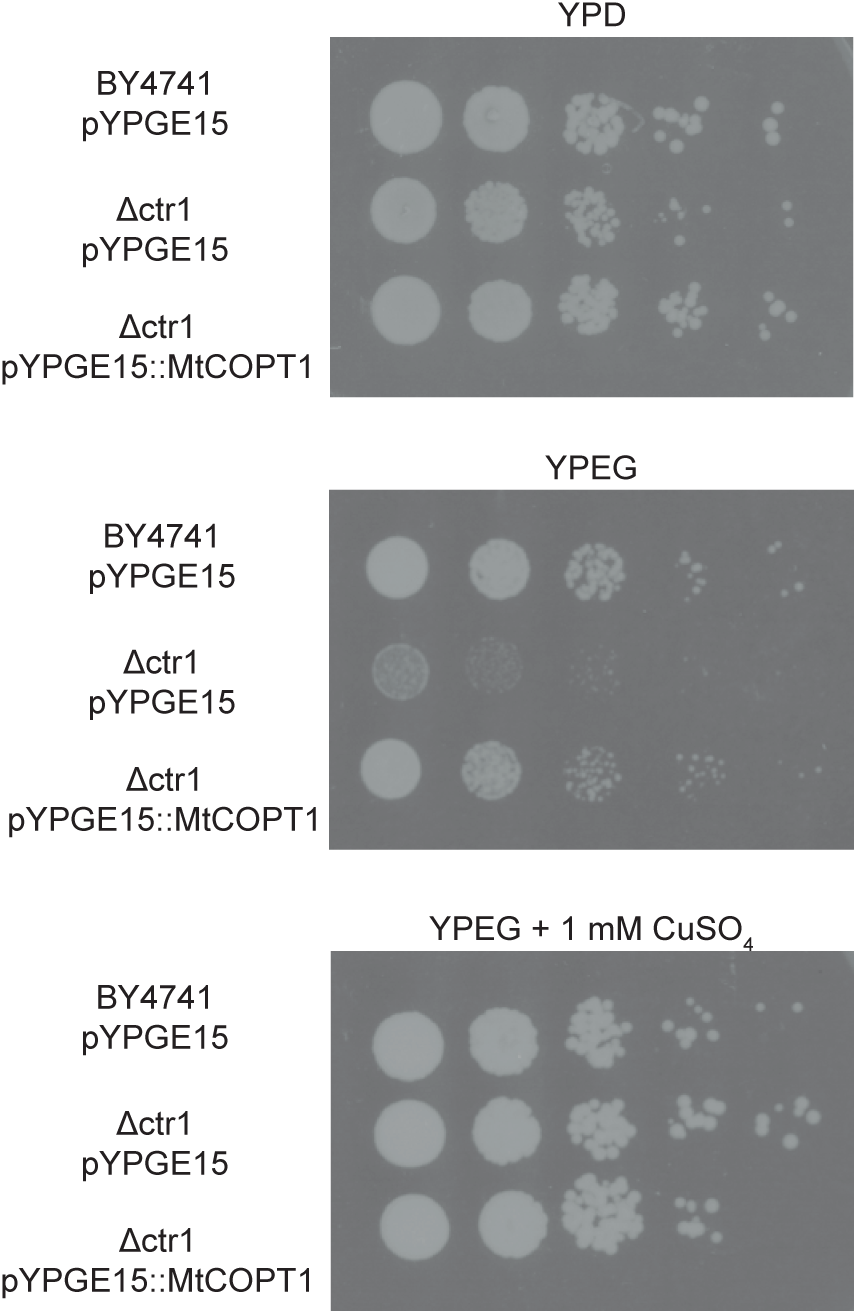
MtCOPT1 transports copper towards the cytosol. Yeast strain BY4741 was transformed with pYPGE15, while BY4741-derived ∆*ctr1* strain was transformed with either empty pYPGE15 or with pYPGE15 containing *MtCOPT1* coding sequence. Serial dilutions (10x) of each transformant were grown for 2 days at 28ºC in non-selective medium (YPD), in fermentative selective YPEG medium and in YPEG supplemented with 1 mM CuSO_4_.

### MtCOPT1 is a plasma membrane protein expressed in the nodule late differentiation, interzone, and early fixation zones

The different developmental zones of an indeterminate type nodule carry out different biological functions (Vasse et al., 1990). Consequently, as a first approach to discern the biological role of MtCOPT1, the expression profile in the nodule was determined by fusing the 2 kb region upstream of the start codon of *MtCOPT1* to a *β‐glucuronidase* (*gus*) gene. Analysis of GUS activity in nodules of *A. rhizogenes*‐transformed roots 28 days post inoculation (dpi) showed that expression was confined to the late infection/differentiation, interzone, and early fixation zone (Fig. 3a and b). Similar results were detected when using a *MtCOPT1*promoter::*green fluorescent protein* (*gfp*) fusion (Supporting Information, Fig. S2). This expression profile was very similar to the one recorded in the Symbimics database (https://iant.toulouse.inra.fr/symbimics/) that shows the transcripts in each nodule region (Fig. 3c) (Roux et al., 2014).

**Figure 3.**
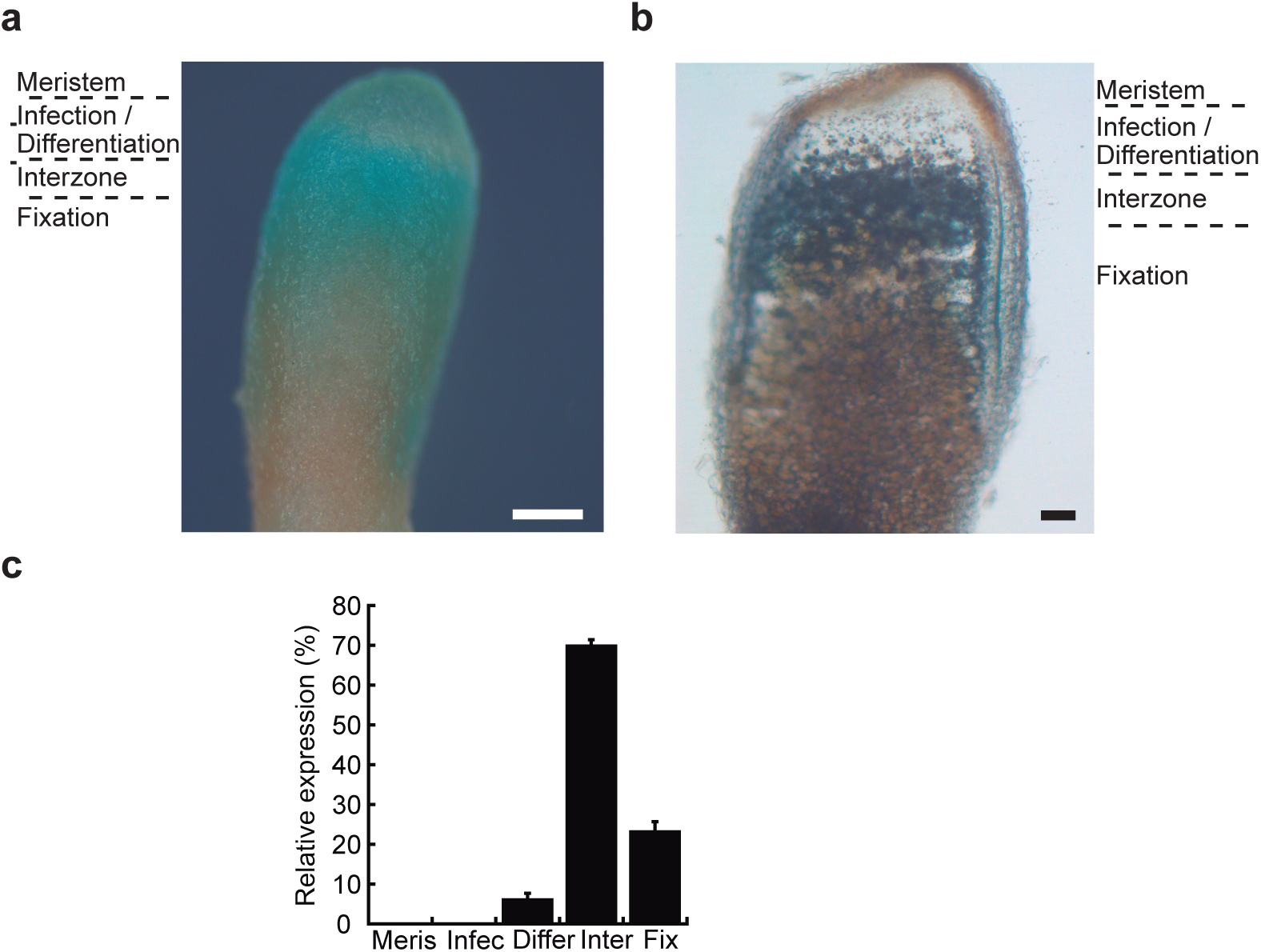
*MtCOPT1*gene is expressed in the infection, interzone and nitrogen fixationzones. (a) GUS staining of *M. truncatula* 28-dpi nodules expressing the *gus* gene under the control of *MtCOPT1* promoter. Bar =300 μm. (b) Longitudinal section of a 28-dpi nodule expressing the *gus* gene under the control of *MtCOPT1* promoter. Bar = 100 μm.(c)Expression of *MtCOPT1* in *M. truncatula* nodules determined by laser-capture microdissection coupled to RNA sequencing. Data were obtained from the Symbimics database (https://iant.toulouse.inra.fr/symbimics/). Meris, meristem; Infec, infection one; Differ, differentiation zone; Inter, interzone; Fix, nitrogen fixation zone.

The above expression profile was also validated by detecting the protein localization using an epitope-labelled MtCOPT1 that has three hemagglutinin (HA) tags fused to its N-terminal region. Expression of this construct was driven by the same promoter region as for the GUS activity visualization studies. MtCOPT1-HA was located in cells in the late infection/differentiation, interzone, and in the younger parts of the fixation zone (Fig. 4a). The protein was detected in both rhizobia-infected and non-infected cells. This result was not due to autofluorescence, since sections that were not incubated with the primary antibody did not show any signal (Supporting Information, Fig. S3). A closer view of these cells showed a peripheral distribution of the protein, indicative of a plasma-membrane localization (Fig. 4b). To validate this subcellular localization, *N. benthamiana* leaves where co-agroinfiltrated with a plasmid expressing a N-terminal GFP-labelled *MtCOPT1* under a 35S promoter and a plasma membrane marker fused to the cyan fluorescent protein (CFP). Confocal imaging of both constructs showed colocalization (Fig. 4c), validating the putative plasma membrane localization of MtCOPT1. No GFP signal was found in cells expressing the CFP-labelled PM marker alone, nor was CFP signal detected in cells containing only MtCOPT1-GFP (Supporting Information, Fig. S4), thus ruling out any non-specific signal when both constructs are co-expressed.

**Figure 4.**
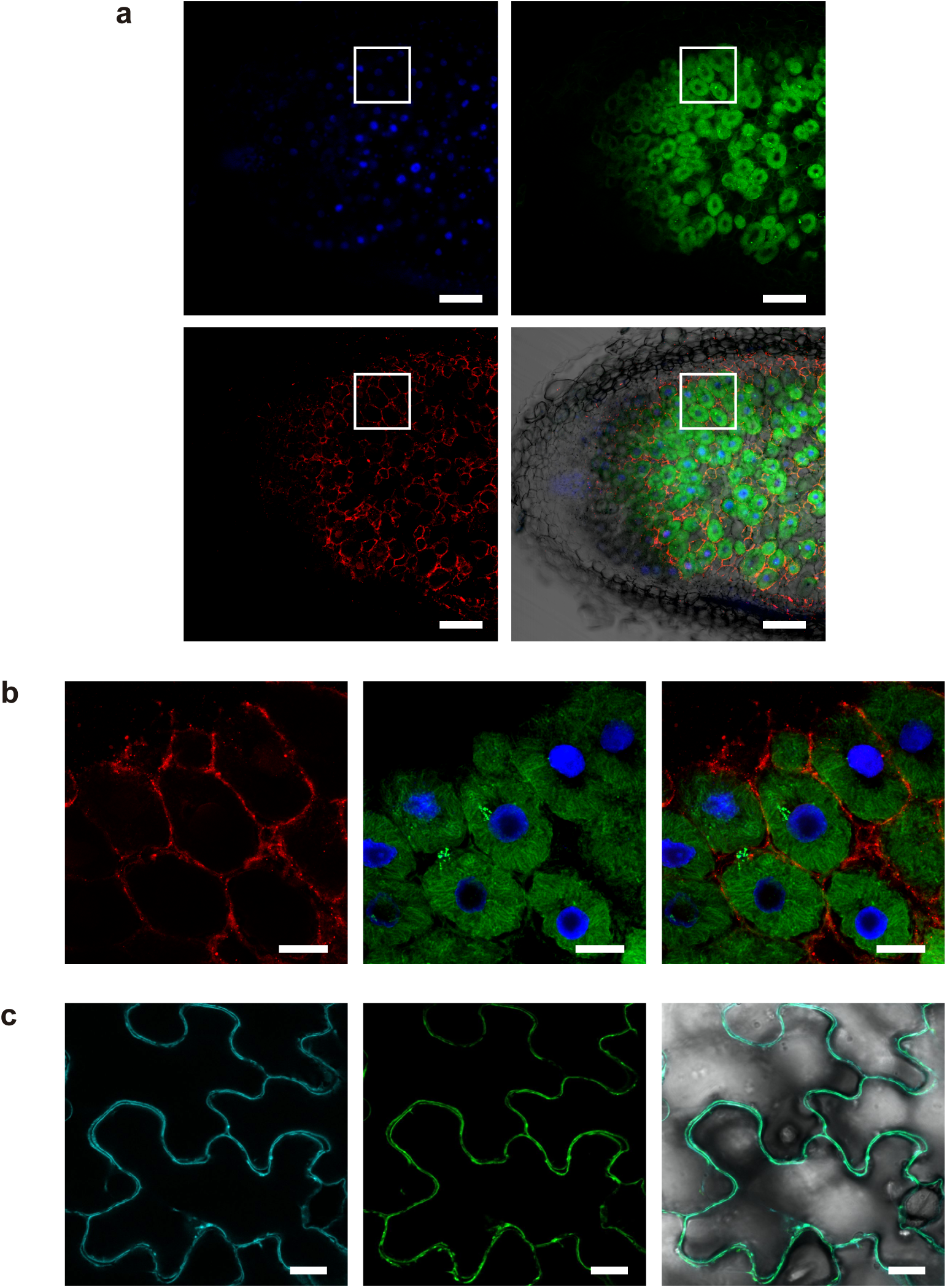
Subcellular localization of MtCOPT1-HA. (a) Cross section of a 28-dpi *M. truncatula* nodule inoculated with *S. meliloti* constitutively expressing GFP (green, upperright panel) and transformed with a vector expressing the fusion *MtCOPT1-HA* under the regulation of its endogenous promoter. Nodules were stained with DAPI to show DNA (blue, upper left panel). MtCOPT1-HA localization was determined using an Alexa 594-conjugated antibody (red, lower left panel). The lower right panel shows the overlay of the transillumination, DNA, *S. meliloti*, and MtCOPT1-HA. Scale bar =100 μm. (b) Detailed view of rhizobia-infected cells. GFP-expressing *S. meliloti* are shown in green, red indicates the position of MtCOPT1-HA, and blue is DAPI-stained DNA. Scale bar = 25 μm. (c) Localization of plasma membrane marker pm-CFP transiently expressed in tobacco leaf cells (left panel) and localization of MtCOPT1-GFP transiently expressed in the same cells (central panel). Right panel shows the overlaid images and the transillumination. Scale bar = 25 μm.

### Loss of *MtCOPT1* function results in impaired, copper-mitigated nitrogenase activity

To determine the role of MtCOPT1 in *M. truncatula,* the *Transposable Element from N.tabacum* (*Tnt1*) mutant line NF19829 (*copt1-1*) was obtained from the Noble Foundation insertion mutant library (Tadege et al., 2008). This line carries a *Tnt1* insertion in position +32, that fully silences *MtCOPT1* expression (Fig. 5a). Mutating *MtCOPT1-1* had no significant effect on the expression levels of the other *M. truncatula COPT* family members (Supporting Information, Fig. S5)

**Figure 5.**
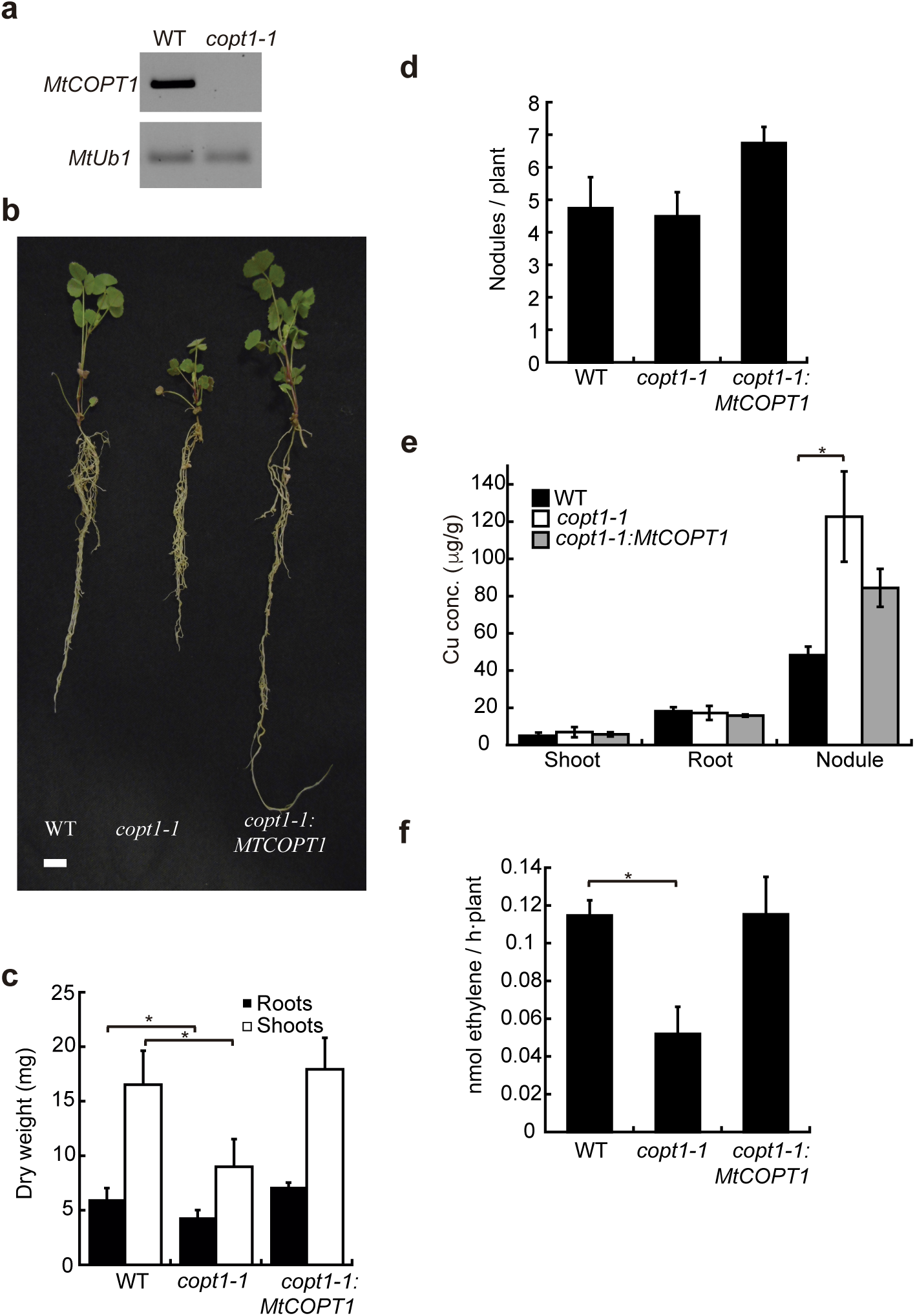
*MtCOPT1* mutation results in a reduced nitrogen fixation rate. (a) RT-PCRamplification of *MtCOPT1* transcript in 28-dpi nodules of *M. truncatula* wild-type (WT) and mutant (*copt1-1*) plants. *Ubiquitin carboxyl-terminal hydrolase1* (MtUb1) was used as control for PCR amplifications. (b) Growth of representative plants of wild type, *copt1-1*, and *copt1-1* transformed with a wild-type copy of *MtCOPT1*. Scale bar = 1 cm. (c)Biomass production in shoots and roots. Data are the mean ± SD of at least 6 independently transformed plants. (d) Number of nodules per plant. Data are the mean ± SD of at least 6 independently transformed plants. (e) Copper content in shoots, roots, and nodules of wild type, *copt1-1*, and *copt1-1* transformed with *MtCOPT1*. Data are the mean ± SD of three sets each with five independently transformed plants. (f) Nitrogenase activity in 28-dpi nodules. Acetylene reduction was measured in duplicate in three sets, each of four independently transformed plants. Data are the mean ± SD. Asterisk indicates significant differences (p<0.05).

Since *MtCOPT1* was expressed solely in nodules, the phenotype of *copt1-1* was assessed in *S. meliloti* inoculated plants watered with a nitrogen-deficient nutritive solution. Under these conditions, *copt1-1* showed reduced growth (Fig. 5b) and biomass (Fig. 5c) compared to wild-type plants. However, no significant differences were found in nodule number (Fig. 5d). Consistent with the role of MtCOPT1 in Cu^+^ transport in nodules, alterations in copper levels were detected in these organs, with over twice as much copper in *copt1-1* nodules than in wild-type ones (Fig. 5e). However, no major change in copper distribution was observed with copper sensor CS1 (Bernal et al., 2012; Chan et al., 2012) (Supporting Information, Fig. S6). Copper levels did not significantly differ between roots of wild-type or *copt1-1* plants, or between shoots of these two genotypes. Acetylene reduction activity (Dilworth, 1966) was measured to determine how nitrogenase activity was affected by mutating *MtCOPT1*. The results showed a *ca.* 50% reduction of this activity in the *copt1-1* mutant (Fig. 5f). This phenotype can be attributed to the *Tnt1* insertion in *MtCOPT1* since it was restored by transforming *copt1-1* with *MtCOPT1* regulated by its own promoter (Fig. 5). To test whether the phenotype observed was caused by altered copper delivery to the plant, a plant nutrient solution fortified with a 100-fold excess of copper compared to our standard was used (Fig. 6). This resulted in improved growth of *copt1-1*, similar to that of wild type plants (Fig. 6a), and a restoration of biomass production (Fig. 6b) and of nitrogenase activity (Fig. 6c). No differences between wild-type and *copt1-1* plants were observed when they were not inoculated and an assimilable nitrogen source was provided in the nutrient solution (Supporting Information, Fig. S7).

**Figure 6.**
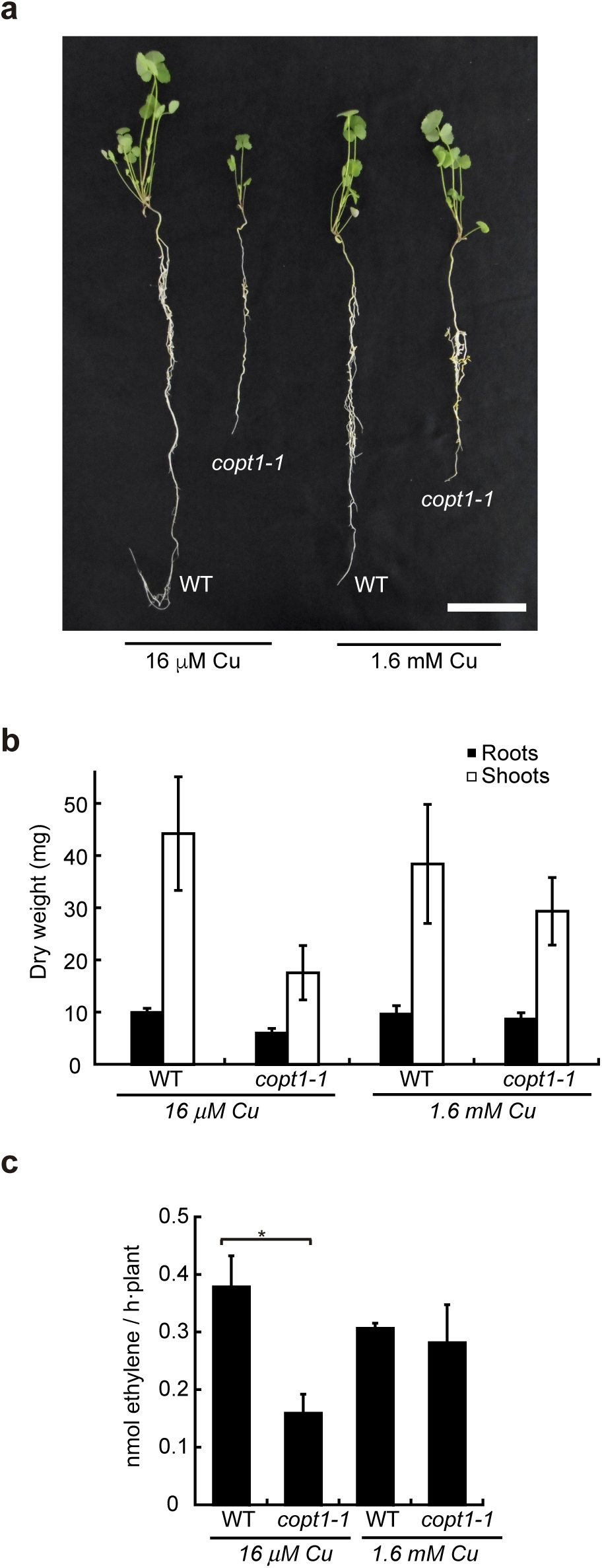
Copper complementation of the *copt1-1* phenotype. (a) Growth of representative plants of wild type, and *copt1-1* watered with standard (16 μM Cu) or copper fortified (1.6 mM Cu) nutrient solution. Scale bar = 3 cm. (b) Biomass production in shoots and roots of wild type, and *copt1-1* watered with standard (16 μM Cu) or copper fortified (1.6 mM Cu) nutrient solution. Data are the mean ± SD of at least 11 plants. (c) Nitrogenase activity in 28-dpi nodules. Acetylene reduction was measured in duplicate in three sets, each of four plants. Data are the mean ± SD. Asterisk indicates significant differences (p<0.05).

### Bacteroid cytochrome oxidase activity is impaired in *copt1-1* nodules

Although reduced nitrogenase activity could cause lower biomass production in *copt1-1* plants, there is no direct link between nitrogenase and copper nutrition. However, this enzymatic activity heavily relies on the obligatorily aerobic energy metabolism of the bacteroid, in which copper-dependent cytochrome oxidase *cbb*_*3*_ plays a critical function. To test whether mutation in *MtCOPT1* had a negative impact on this metabolic process, cytochrome oxidase activity was measured in bacteroids isolated from *copt1-1* and wild-type nodules (Fig. 7). Bacteroids from *copt1-1* had 60 % less activity than the controls. Increasing copper concentration in the nutrient solution restored activity to levels similar to those of the wild type. This phenotype was not the result of down-regulation of the rhizobial cytochrome oxidase-encoding genes, since bacteroids from *copt1-1* had increased expression levels of their two *fixN* genes (Supporting Information, Fig. S8).

**Figure 7.**
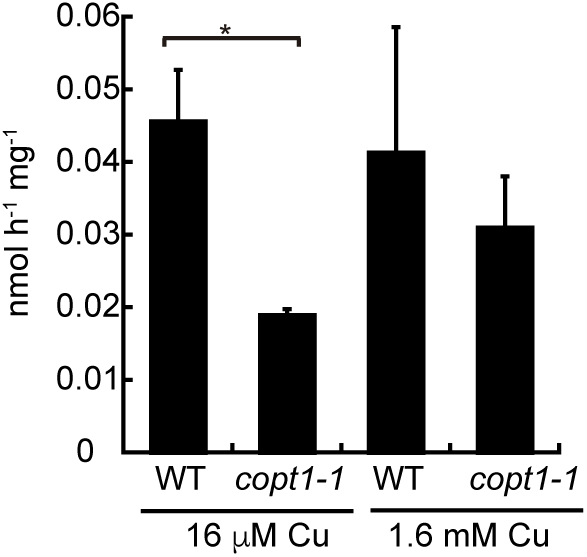
Cytochrome oxidase activity in bacteroids isolated from wild-type and *copt1-1* plants watered with standard (16 μM Cu) or copper fortified (1.6 mM Cu) nutrientsolution. Data are the mean ± SD of three sets of five pooled plants. Asterisk indicates significant differences (p<0.05).

## Discussion

Symbiotic nitrogen fixation heavily relies on a number of metalloproteins to carry out this complex and energetically costly reaction (Brear et al., 2013; González-Guerrero et al., 2014; González-Guerrero et al., 2016). Therefore, studying how metals are allocated from the host plant to the nitrogen-fixing rhizobia is of great importance in view of renewed efforts to engineer nitrogen fixation capabilities in non-legumes (Oldroyd & Dixon, 2014; Ivleva et al., 2016; Lopez-Torrejon et al., 2016; Mus et al., 2016). In this context, a substantial effort has been dedicated to studying how iron is delivered to the nodule and released into the apoplast (Rodríguez-Haas et al., 2013), a process likely facilitated by citrate (Takanashi et al., 2013), to identify the plant transporters involved in iron transport in rhizobia-infected cells (Kaiser et al., 2003; Hakoyama et al., 2012; Tejada-Jiménez et al., 2015), and to describing mechanisms of iron buffering in the bacteroid (Zielazinski et al., 2013). However, less is known about other transition metals required for critical functions in symbiotic nitrogen fixation.

Copper is involved in several plant physiological processes: energy transduction (cytochrome oxidase) (Brunori et al., 2005); cell wall metabolism (laccases) (Hakulinen and Rouvinen, 2015); free radical metabolism (superoxide dismutase) (Fridovich, 1976); hormone metabolism (ethylene receptor) (Rodríguez et al., 1999); and cofactor biosynthesis (Cnx1) (Kuper et al., 2004). Copper deficiencies cause severe growth defects in plants, associated with reduced photosynthetic rates and cell wall production (Burkhead et al., 2009). It is also detrimental for symbiotic nitrogen fixation, a process likely associated with reduced cytochrome oxidase activity in the bacteroids (O'Hara, 2001). Bacteroids carry out aerobic metabolism at extremely low free oxygen concentrations in nodules, which requires the high-affinity cytochrome oxidase *cbb*_*3*_ to satisfy the energy demands of nitrogenase (Preisig et al., 1996b). This iron-copper enzyme is assembled from the *fixNOQP* operon, but the copper cofactor is provided by a subset of Cu^+^-ATPases (FixI in rhizobia) that extrude this ion from the bacteroid cytosol (Kahn et al., 1989; Preisig et al., 1996a; Raimunda et al., 2011; Patel et al., 2014s). Both *fixNOQP* and *fixGHIS* are expressed only in bacteroids, indicating roles in adaptation to the endosymbiotic lifestyle. While we know how bacteroids transfer copper to this cytochrome oxidase, less is known about how the metal is delivered by the host plant. In analogy to how iron is delivered (Rodríguez-Haas et al., 2013; Tejada-Jiménez et al., 2015), it might be hypothesized that it is carried out by the vasculature and released in the apoplast of the infection/differentiation zone. Following this, a plasma membrane copper transporter would introduce this element into the cell to be then delivered across the symbiosome membrane to the bacteroids. Our results indicate that MtCOPT1 is responsible for this apoplastic copper uptake.

*MtCOPT1* is a copper transporter expressed only in nodules. Yeast complementation assays indicate that it is involved in copper uptake. *MtCOPT1*-promoter∷*gus/gfp* fusion studies indicate that this role is carried out in the region from the late infection/differentiation zone to the early fixation zone, where it has been suggested that copper would be released from the vasculature into the nodule apoplast (Rodriguez-Haas et al., 2013). RNAseq data reported in the Symbimics database and obtained from laser-captured microdissected cells in this region validate our expression results (Roux et al., 2014). Moreover, the putative role of MtCOPT1 in introducing copper into nodule cells is supported by the localization of HA-tagged MtCOPT1 in the periphery, very likely the plasma membrane, of infected and non-infected cells. This was confirmed by colocalization with a plasma membrane marker in tobacco leaves.

Normal plant growth under symbiotic conditions is dependent on MtCOPT1 activity, indicated by the reduced biomass production of *copt1-1* plants when compared to the wild-type. This was the result of a *ca.* 50% reduction of nitrogenase activity, and not to alterations in nodule development. This phenotype was caused by the *Tnt1* insertion in *MtCOPT1* and not by any other insertion in a different part of the genome, since transformation with *MtCOPT1* expressed under the control its own promoter was able to restore wild-type growth and nitrogenase activity in *copt1-1*. However, MtCOPT1 transports Cu^+^, and there is no evidence for a COPT/Ctr transporter with the ability to transport iron or molybdenum, the two metal cofactors directly involved in nitrogenase catalytic mechanism (Miller et al., 1993; Rubio and Ludden, 2005). Therefore, MtCOPT must support nitrogen fixation indirectly, via a copper-dependent process. This idea is supported by restoration of the wild-type phenotype after watering plants with a copper-fortified nutrient solution.

One of the processes likely affected by alterations in copper homeostasis in the nodule is cytochrome oxidase *cbb*_*3*_ activity in the bacteroids. A malfunction in this enzyme could result in a decrease in energy metabolism and/or an increase in free-oxygen concentration, either of which could negatively affect nitrogenase activity. In fact, a mutation in the *Bradyrhizobium japonicum fixNOQP* operon results in a *fix*^*¯*^ phenotype (Preisig et al., 1996b). A similar phenotype can be observed merely by mutating *fixI*, the P_1b_-ATPase that provides copper for this enzyme (Preisig et al., 1996a; Patel et al., 2014). Bacteroids isolated from *copt1-1* nodules also showed a significant reduction in cytochrome oxidase activity, consistent with a decrease in copper supply to bacteroids. However, the *copt1-1* phenotype did not appear to be as severe as that of mutants in *fixI* or in *fixN*, since some nitrogenase activity remained in the mutant nodules and no major developmental change was observed. This indicates that MtCOPT1 is not the only transporter responsible for copper uptake by these cells. The close homologue, MtCOPT2 is an unlikely candidate for this role, since its expression cannot be detected in nodules. On the other hand, MtCOPT3, the only other COPT family member expressed in nodules, could conceivably carry out this role. However, its expression is not affected by *MtCOPT1* mutation, which suggests that its role is independent of MtCOPT1, while its expression in every plant organ is indicative of a more general role in the plant physiology. Alternatively, another family of metal transporters, such as YSL, ZIP or Nramp, could carry out this role.

Our results also indicate the existence of a systemic metal deficiency signal originating in nodule cells. We detected a two-fold increase in copper levels in *copt1-1* mutant nodules compared to those of the wild type, which was somewhat of a surprise given the postulated role of MtCOPT1 in copper uptake by nodule cells. A possible explanation for this result is that cellular demand for copper in nodules is signalled systemically to increase supply to nodules until demand is met. In the absence of MtCOPT1 activity, intracellular levels of copper remain low, possibly triggering a systemic signal(s) that results in more copper being transported into nodules. Increased copper concentration in the apoplast appears to be sufficient to allow alternative transporters (e.g. COPTs, NRAMPs) to import some copper into cells, which would explain why some cytochrome oxidase activity was still detected in *copt1-1* nodules. However, the affinity for copper of alternative transporters must be relatively low, because very high concentrations of copper (100x) in the nutrient solution were required to complement the mutant phenotype. Similar observations have been made for the nodule-specific molybdate transporter *MtMOT1.3* (unpublished).

In conclusion, copper entering nodules is very likely delivered by the vasculature and released in the infection zone-interzone where MtCOPT1 would transport it into cells (Fig. 8). Within the cytoplasm, a Cu^+^-chaperone (Robinson & Winge, 2010) would probably deliver the cation to other transporters and apo-proteins. In the case of the symbiosome, it could be hypothesized that a P_1b_-ATPase mediates copper delivery to the bacteroid, since these ATPases have been shown to mediate copper transport (Burkhead et al., 2009; Kaplan & Lutsenko, 2009). However, no nodule-specific or nodule-induced Cu^+^-ATPase has been reported to date either experimentally or from the available transcriptomic databases. Once in the symbiosome space, copper would be transported into the bacteroid, via transporters in both the outer and inner bacteroid membranes, and subsequently delivered to cytochrome oxidase *cbb*_*3*_ via FixI (Preisig et al., 1996a; Patel et al., 2014; Trasnea et al., 2016; Fig. 8). This model does not exclude the existence of other cuproproteins affected by *MtCOPT1* mutation, since once copper is within the cytosol, it can be used in several different ways, and some of them could also potentially affect nitrogenase activity in nodules.

**Figure. 8.**
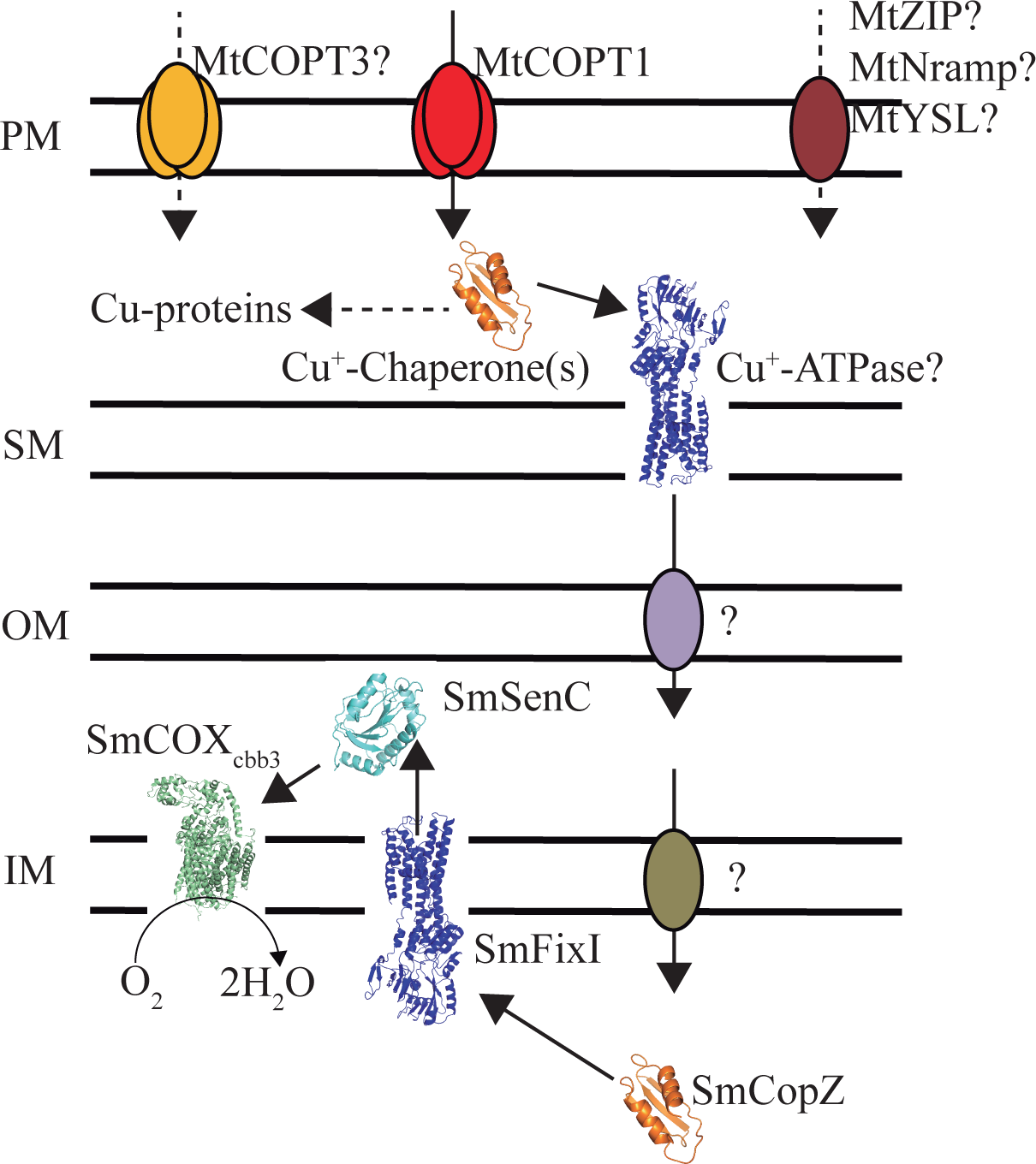
Model of copper homeostasis in rhizobia-infected nodule cells. Copper is introduced into the host cell cytosol by MtCOPT1. There, Cu^+^ will be transferred by cytosolic Cu^+^-chaperones to other cuproproteins or transported across the symbiosome membrane, very likely by Cu^+^-ATPases. Copper through some unknown transporters is delivered into the bacteroid cytosol, where it will be bound by Cu^+^-chaperone CopZ. This protein will transfer copper to the Cu^+^-ATPase FixI. In the periplasm, copper will be delivered to SenC, which will add the copper cofactor to cytochrome oxidase *cbb*_*3*_. In addition to MtCOPT1, other copper uptake systems must exist. Candidates are MtCOPT3, the other COPT family member expressed in nodules, or a member of the Nramp, ZIP, or YSL metal transport families. PM stands for plasma membrane, SM for symbiosome membrane, OM for bacteroid outer membrane, and IM for bacteroid inner membrane.

## Acknowledgments

This research was funded by the Spanish Ministry of Economy and Competitiveness (grant number AGL-2012-32974) and by the European Research Council Starting Grant (grant number ERC-2013-StG-335284) to MG-G. RC-R was supported by a Formación del Personal Investigador fellowship (BES-2013-062674). Part of the work was funded by the US National Science Foundation Plant Genome Research Program (grant IOS1127155 to MU). The authors would like to thank Dr. Chris Chang for providing the CS1 sensor, and Dr. José M. Argüello for sending us the DsRed-expressing *S. meliloti.*

### Author contribution

Phylogenetic tree, protein secondary structure prediction, yeast complementation, and promoter:*gus* studies were carried out by M.S. Gene expression was determined by M.S., and R.C.-R, as well as confocal microscopy studies. The *copt1-1* mutant was obtained by I.K., and M.K.U. Phenotypic characterization of *copt1-1* was performed by M.S., R.C.-R., I.A., and V.E. Bacteroid cytochrome oxidase activity was determined by I.A., and M.S. J.I. and M.G.-G. were responsible for experimental design, data analyses, and wrote the manuscript with contributions from all the authors.

